# High throughput analysis of vacuolar acidification

**DOI:** 10.1101/2022.08.13.503847

**Authors:** Chi Zhang, Adam Balutowski, Yilin Feng, Jorge D. Calderin, Rutilio A. Fratti

**Affiliations:** Department of Biochemistry, University of Illinois Urbana-Champaign, Urbana IL, 61801, USA; Center for Biophysics and Quantitative Biology, University of Illinois Urbana-Champaign, Urbana IL, 61801, USA; Biochemistry, Biophysics, and Structural Biology program, Washington University, St. Louis, MO

**Keywords:** V-ATPase, Vph1, Lysosome, Acridine Orange, Lanthanum, Gadolinium, Nickel

## Abstract

Eukaryotic cells are compartmentalized into membrane-bound organelles, allowing each organelle to maintain the specialized conditions needed for their specific functions. One of the features that change between organelles is luminal pH. In the endocytic and secretory pathways, luminal pH is controlled by isoforms and concentration of the vacuolar-type H^+^-ATPase (V-ATPase). In the endolysosomal pathway, copies of complete V-ATPase complexes accumulate as membranes mature from early endosomes to late endosomes and lysosomes. Thus, each compartment becomes more acidic as maturation proceeds. Lysosome acidification is essential for the breakdown of macromolecules delivered from endosomes as well as cargo from different autophagic pathways, and dysregulation of this process is linked to various diseases. Thus, it is important to understand the regulation of the V-ATPase. Here we describe a high-throughput method for screening inhibitors/activators of V-ATPase activity using Acridine Orange (AO) as a fluorescent reporter for acidified yeast vacuolar lysosomes. Through this method, the acidification of purified vacuoles can be measured in real-time in half-volume 96-well plates or a larger 384-well format. This not only reduces the cost of expensive low abundance reagents, but it drastically reduces the time needed to measure individual conditions in large volume cuvettes.

## 1. Introduction

Eukaryotic cells are subdivided and organized into membrane-bound organelles that have distinct sets of enzymes and molecules to serve in organelle-specific roles [1,2]. Organelles communicate through the budding, trafficking, and fusion of transport vesicles containing luminal or membrane integrated cargo. In the endolysosomal pathway, cargo is transferred from early endosomes to late endosomes and ultimately lysosomes. Upon delivery to lysosomes macromolecular cargo is broken down into building blocks by various digestive enzymes, which are activated by the acidification of the lysosomal lumen [2–6].

One of the major characteristics that distinguishes early endosomes from late endosomes and lysosomes is the gradual decrease of pH. As endosomes mature, the luminal pH drops from 6.5 in early endosomes to 5.5 in late endosomes, while lysosomal compartments acidify to a range between 4 and 5, depending on the organism [2,6–8]. Luminal acidification is required in the activation of and function of lysosomal digestive enzymes to breakdown proteins, lipids, toxins as endosomal cargo, and entire organelles delivered to lysosomes through autophagy [2,6–9]. The principal source of vesicle acidification is vacuolar-type H^+^-ATPase (V-ATPase) that is composed of two protein sub-complexes. The V_O_ sub-complex is imbedded in the membrane and contains the C-ring that binds and releases H^+^. As it rotates the C-ring encounters the two hemichannels of the A-subunit that face the cytosol and vesicle lumen. The cytosolic hemi-channel loads the H^+^ on to the C-ring and the luminal hemi-channel releases the H^+^ into the vesicle lumen. The energy used to transfer H^+^ through the V_O_ against a concentration gradient comes from the cytoplasmic V_1_ complex containing the ATPase activity of the enzyme. When assembled, the V_1_-V_O_ holocomplex uses ATP hydrolysis to turn the C-ring and pump H^+^ into the lysosomal lumen [2,10,11]. The importance of lysosomal acidification is broad. Not only can defective acidification lead to the accumulation of immature hydrolytic enzymes, but deficient V-ATPase activity has been linked to reduced autophagy, lysosomal storage disorders, Alzheimer’s Disease, Parkinson’s Disease, and more [8,10,12,13].

Since the process of endosome maturation and endocytosis is evolutionarily conserved in eukaryotic cells, it is advantageous to study the lysosomal fusion pathway and V-ATPase function in the budding yeast *Saccharomyces cerevisiae*. One of the advantages of using *S. cerevisiae* vacuolar lysosomes is the ease of their isolation at high concentrations. Vacuoles isolated from one liter of medium can yield upwards of 1 mg of vacuoles by protein. Importantly, these organelles contain all the essential proteins and lipids for *in vitro* analysis of fusion, fission, Ca^2+^ transport, actin polymerization and V-ATPase activity [11,14–18]. Furthermore, the relative ease and low cost of yeast genetics makes it easy to introduce gene deletions, mutations, truncations, and chimeras to further probe cellular functions including vacuole acidification [19,20]. A common method to quantify yeast vacuole acidification uses the fluorescein derivative BCECF-AM (2′,7′ – bis-(Carboxyethyl)-5-(and-6)-carboxyfluorescein-acetoxymethyl ester), a pH-sensitive fluorescent probe that labels mammalian cytosol and accumulates in yeast vacuoles. BCECF fluoresces at two excitation wavelengths. While excitation at 440 nm is not affected by pH while pH does affect excitation at 490 nm. Ratios of emission at 535 nm are taken with each excitation point to calculate the pH of vacuole lumen [21–24]. Other dyes, including Oregon Green, fluorescein derivatives, and HPTS (8-hydroxypyrene-1,2,3-trisulfonic acid), pHrodo, as well as the pH sensitive GFP derivative pHlourin have been used in fluorescently labeling lysosomes of mammalian cells [25–33]. These assays, however, measures the lysosome pH *in vivo*, which is not ideal for highthroughput studies of V-ATPase function. This is due to the need to feed cells dextran conjugates of pH sensitive dyes or genetic expression of pHlouorin. For our high-throughput *in vitro* assay we used the fluorescent dye Acridine Orange (AO) which readily accumulates in acidified isolated vacuoles in vitro [34–36].

AO has two dominant absorption peaks at 460 nm and 540 nm and the difference in absorbance has been used as a measure of vacuole acidification [34,37]. Using this difference in absorbance, others have used AO to measure in vitro yeast vacuole acidification, but this can be laborious and can limit the number of variables tested per vacuole preparation. However, instead of relying on a difference in absorbance, we take advantage of changes in fluorescence intensity. AO fluoresces with a peak emission at 530 nm when excited at 488 nm [38,39] and stains double-stranded DNA with the expected green color, but stains single-stranded DNA and RNA with red color [40–43]. The reason for the two AO emission wavelengths is because of its metachromatic shift. When AO is accumulated, the increase of local concentration of AO pushes monomers to dimerize through base stacking, and the dimerization leads to a shift in dominant emission peak from 530 nm to 680 nm [39,40,44,45]. When AO is used to stain acidic compartments *in vivo*, AO monomers pass through membranes and become trapped in acidic compartments upon protonation to fluorescently label these compartments [43,44].

Here we present a high-throughput *in vitro* assay using AO fluorescence and half-volume 96 well plates that can be scaled to a 384-well format. Upon the addition of ATP, the V-ATPase transports H^+^ from the reaction buffer system to the vacuole lumen and changes in fluorescence are measured with an excitation of 485 nm and emission at 520 nm. As V-ATPase complexes hydrolyze ATP to pump H^+^ into the vacuole lumen, AO accumulates in the acidic vacuoles, which causes to a shift in emission wavelength from 520 nm to 680 nm due to AO dimerization [38,39,44], therefore a gradual decrease of fluorescence signals at emission wavelength of 520 nm is observed. Thus, a larger decrease of AO fluorescence indicates greater vacuolar acidification.

## 2. Experimental

### 2.1. Chemicals/Reagents

Soluble reagents were dissolved in PIPES-Sorbitol (PS) buffer (20 mM PIPES-KOH, pH 6.8, 200 mM sorbitol) with 125 mM KCl unless indicated otherwise. ATP was purchased from RPI (Mount Prospect, IL). Bafilomycin A1 and Concanamycin A were from Cayman Chemical (Ann Arbor, MI) and dissolved in DMSO. Acridine orange, Coenzyme A (CoA), Creatine kinase, and FCCP (Carbonyl cyanide-4-(trifluoromethoxy) phenylhydrazone) were purchased from Sigma (St. Louis, MO). Creatine phosphate was from Abcam (Waltham, MA). Pbi2 (Proteinase B inhibitor 2) was prepared as described and dialyzed against PS buffer with 125 mM KCl. [46].

### 2.2. Yeast strains

Vacuoles from BJ3505 genetic backgrounds were used for H^+^ transport assays. *VPH1* was deleted by homologous recombination using PCR products amplified from the pAG32 plasmid with primers 5’-VPH1-KO (5’ – GCTTAGAGGGCTACCTGTGTGTATTTGCATGGGTAAAAA-GCCTGAACTCACC – 3’) and 3’-VPH1-KO (5’ – AGTCCTCAAAATTTAGCTTGAAGCGGTTATTCCTTTGCCCTCGGACGAGTG – 3) with homology flanking the *VPH1* coding sequence. The PCR product was transformed into chemically competent yeast by standard lithium acetate methods and plated on YPD containing Hygromycin (200 µg/ml) to generate BJ3505 *vph1::hphMX4*.

### 2.3. Vacuole isolation and vacuole acidification

Vacuoles were isolated as described [47]. The proton pumping activity of isolated vacuoles was performed as described by others with modifications [48,49]. *In vitro* H^+^ transport reactions (60 µl) contained 20 µg vacuoles from BJ3505 backgrounds, reaction buffer (20 mM PIPES-KOH pH 6.8, 200 mM sorbitol, 125 mM KCl, 5 mM MgCl_2_), ATP regenerating system (1 mM ATP, 0.1 mg/ml creatine kinase, 29 mM creatine phosphate), 10 µM CoA, 283 nM Pbi2, and 15 µM of the fluorescent dye acridine orange. Reaction mixtures were loaded into a black, half-volume 96-well flat-bottom plate with nonbinding surface (Corning). ATP regenerating system or buffer was added, and reactions were incubated at 27°C while acridine orange fluorescence was monitored. Samples were analyzed in a fluorescence plate reader (POLARstar Omega, BMG Labtech) with the excitation filter at 485 nm and emission filter at 520 nm. Reactions were initiated with the addition of ATP regenerating system 20 sec after the initial measurement. After fluorescence quenching plateaus were reached, we added 30 µM FCCP to collapse the proton gradient and restore acridine orange fluorescence.

### 2.4. Statistical analysis

Results are expressed as the mean ± S.E. Experimental replicates (n) is defined as the number of separate experiments. Significant differences were calculated using One-Way ANOVA for multiple comparisons using Prism 9 (GraphPad, San Diego, CA). *P* values ≤0.05 were considered significant.

## 3. Results and Discussion

To develop a high-throughput assay for V-ATPase function we first set parameters to detect vacuole acidification in an ATP-dependent manner in the absence of cytosol. For this we isolated vacuoles from exponentially growing yeast and used conditions that favor vacuole fusion [34,49]. Upon addition of ATP regenerating system, a rapid drop in AO fluorescence was measured that plateaued after 600 sec **(Fig. 1A)**. In contrast, the reaction lacking ATP did not change in fluorescence intensity, indicating that the observed signal was ATP-dependent. After 600 sec had elapsed, 30 µM FCCP was added to each reaction to equilibrate H^+^ levels across the vacuole membrane and restore starting fluorescence at 544 nm [50]. This shows that the fluorescence decrease at the beginning of the assay is due to accumulation of AO into lysosomes followed by its dimerization.

**Figure 1.**
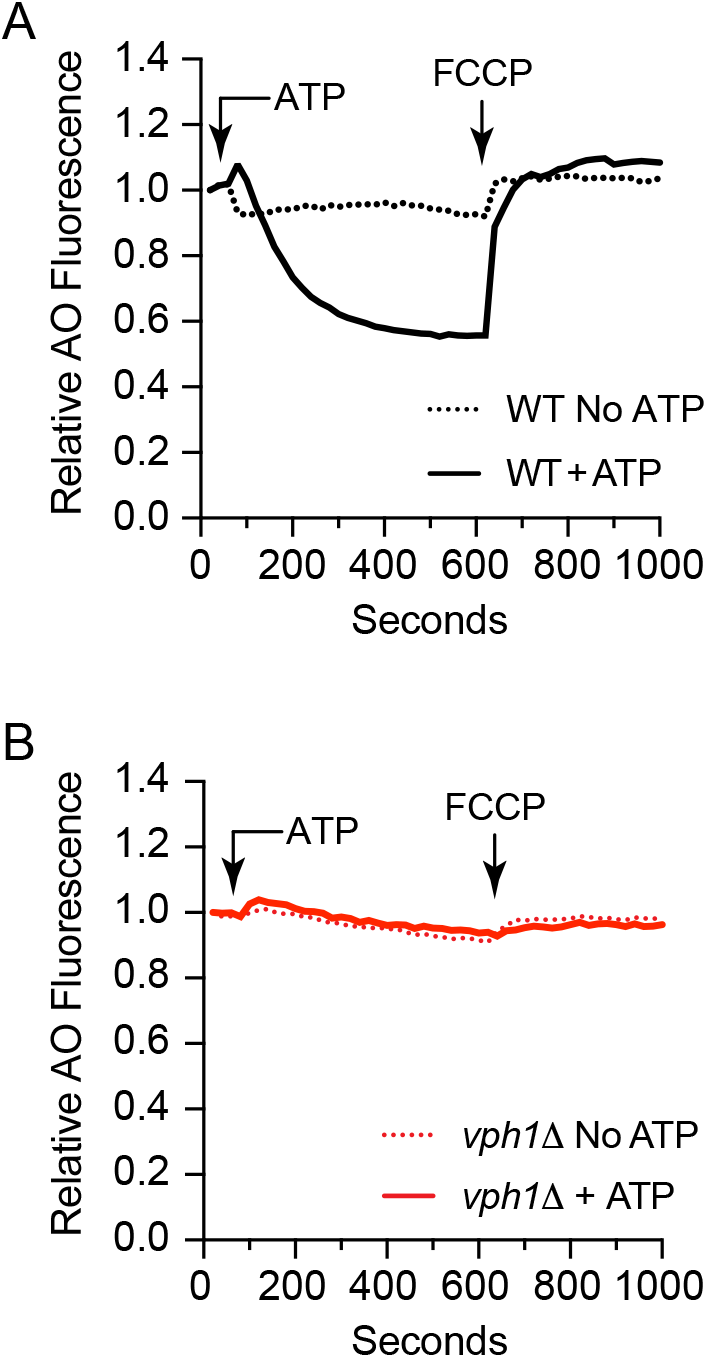
ATP dependent *in vitro* vacuole acidification. Vacuoles were isolated from wild type **(A)** and *vph1*Δ **(B)** yeast and used for proton pumping activity measured by the loss of AO fluorescence at 520 nm. Reactions were incubated with or without ATP regenerating system added at 30 sec and the reactions were incubated for 600 sec. After ∼600 sec reactions were supplemented with 30 µM FCCP was added to collapse the proton gradient and restore AO fluorescence.

Next, we verified whether the decrease in AO fluorescence was dependent on V-ATPase function. To determine this, we used vacuoles isolated from yeast where the V_O_ subunit *VPH1* had been deleted. Others have shown that the Vph1 isoform Stv1 (from the secretory path V-ATPase) can replace Vph1 in the endolysosomal population of the V-ATPase. However, Stv1 does not support vacuolar acidification unless it is overexpressed. This is likely due to the differential regulation of Stv1 and Vph1 by the lipids phosphatidylinositol 4-phosphate (PI4P) and PI(3,5)P_2_, respectively [51,52]. Here we see that *vph1*Δ vacuoles were unable to acidify in the presence or absence of ATP, indicating that the drop in AO fluorescence was due to V-ATPase function **(Fig. 1B)**.

To further illustrate that V-ATPase function was responsible for the ATP-dependent drop in AO signal, we used two well-characterized inhibitors of the holocomplex. Using wild type vacuoles we tested the effects of Bafilomycin A1 and Concanamycin A that block H^+^ translocation by binding to the C-ring of the V_O_ complex [53,54]. Because both inhibitors are dissolved in DMSO we also tested the solvent to see if any observed effects were due to DMSO and not the solutes. In **Figure 2** we show that both Bafilomycin A1 and Concanamycin A fully blocked the drop in AO fluorescence, while DMSO had no effect. These results, along with the data from the *vph1*Δ vacuoles show that the V-ATPase was indeed the reason for the drop in AO fluorescence due to vacuole acidification. Next, we tested if lowering the pH of the vacuole lumen would block AO from shifting in fluorescence. To do this we used the weak base chloroquine, which accumulates in acidic compartments as it is protonated [55,56]. The result is that the luminal pH increases, which would result in a decrease in AO protonation and trapping in the lysosome. This would in turn fail to raise the local concentration of AO in the lysosomes to promote its dimerization and subsequent emission shift. Here we show that AO fluorescence increased as chloroquine concentrations were raised **(Fig. 3)**. This also shows that AO fluorescence can gradually decrease in a dose dependent manner.

**Figure 2.**
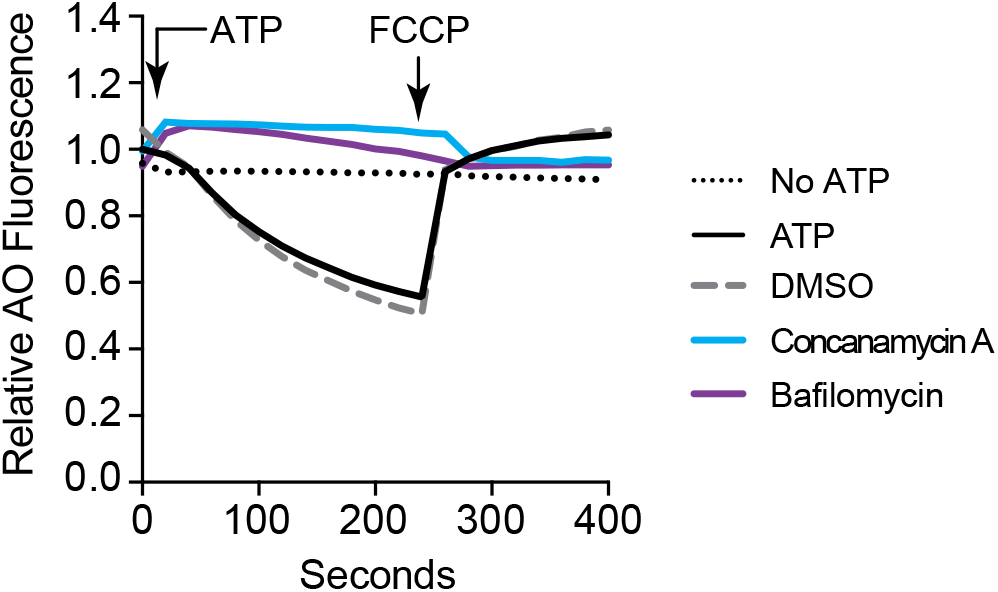
Bafilomycin A1 and Concanamycin prevent vacuole acidification, Vacuoles were isolated from wild type yeast and used for proton pumping activity measured by the loss of AO as described in Figure 1. Reactions were incubated with or without ATP. Reactions containing ATP were treated with PS buffer, 50 µM Bafilomycin, 50 µM Concanamycin or an equivalent of solvent (DMSO). Reactions were incubated for 250 sec before adding FCCP. Readings continued for 400 sec.

**Figure 3.**
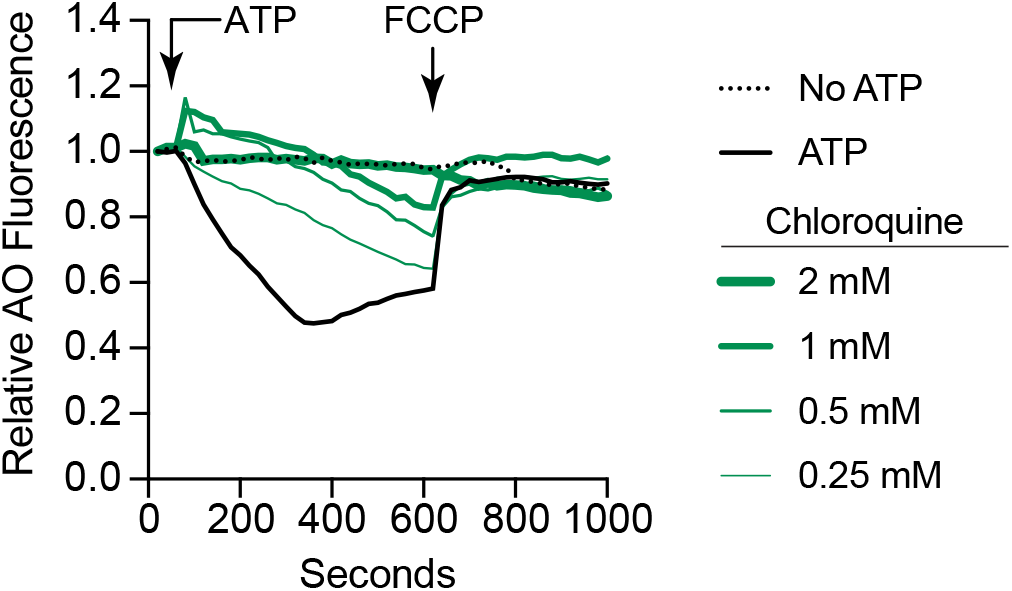
Chloroquine prevents the loss of AO fluorescence. Vacuoles were isolated from wild type yeast and used for proton pumping activity measured by the loss of AO as described in Figure 1. Reactions were incubated with or without ATP. Reactions containing ATP were treated with PS buffer or chloroquine at the indicated concentrations. Reactions were incubated for 600 sec before adding FCCP. Readings continued for 1000 sec.

Finally, we used the high-throughput AO assay to screen for novel inhibitors of V-ATPase function. Previously we found that 100 µM Cu^2+^ potently inhibited vacuole acidification, whereas other divalent cations tested had no effect [49]. Here we tested various metals simultaneously to show how assay can accommodate numerous conditions in a single reading. We tested the lanthanide cations La^3+^ and Gd^3+^ as well as the divalent cation Ni^2+^. Lanthanides were used because their effects on surface charge, dipole potential and transmembrane potential [57]. For instance, Gd^3+^ has been shown to interact with phosphatidylserine and augment dipole potential resulting in increased membrane tension due to enhanced packing of lipids. This can subsequently inhibit the function of mechanosensitive channels including the mucolipin TRPML (Transient Receptor Potential) [58]. That said, both La^3+^ and Gd^3+^ can activate or potentiate the function of other TRP family members including TRPV1 [59]. In addition to channels, lanthanides can also block Ca^2+^-ATPase pumps [60,61]. This is of particular importance since V-ATPase function depends on a Ca^2+^ gradient across the lysosomal membrane [62]. We found that after the addition of ATP the LaCl_3_ and GdCl_3_ treated wells showed an increase in AO fluorescence in a dose-dependent manner. This was not due to vacuole lysis as fluorescence was restored to baseline when H^+^ was equilibrated across the vacuole membrane with the protonophore FCCP **(Fig. 4A-D)**. In contrast to the lanthanides, divalent Ni^2+^ inhibited vacuole acidification in a manner similar to what was seen with Cu^2+^ **(Fig. 4E-F)**. This is also consistent with a previous study showing that Ni^2+^ reduced proton pumping by isolated rat liver lysosomes loaded with fluorescein dextran [63].

**Figure 4.**
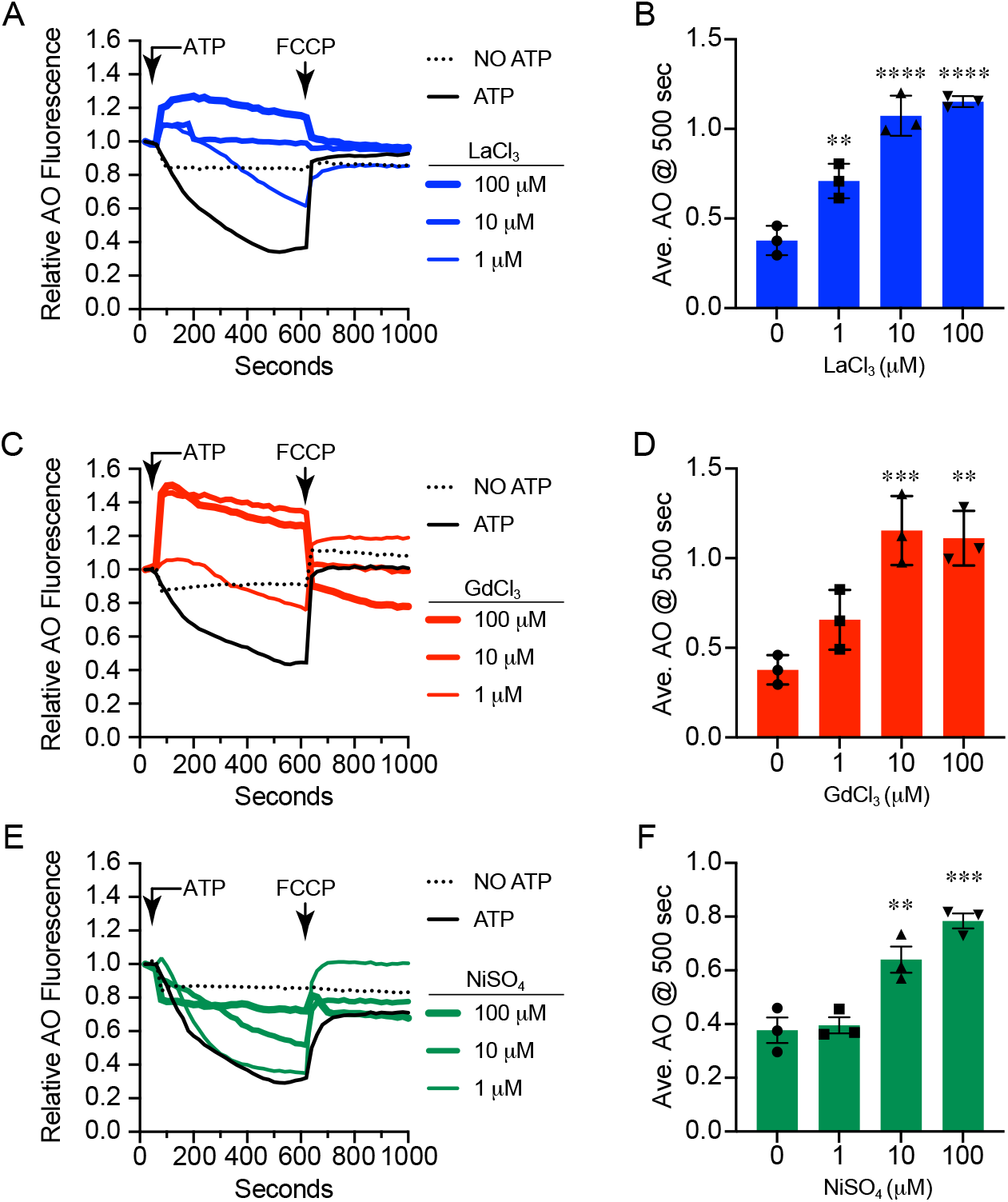
Effect of metals on AO fluorescence. Vacuoles were isolated from wild type yeast and used for proton pumping activity measured by the loss of AO as described in Figure 1. Reactions were incubated with or without ATP. Reactions containing ATP were treated with PS buffer or LaCl_3_ **(A-B)**, GdCl_3_ **(C-D)** or NiSO_4_ **(E-F)** at the indicated concentrations. Reactions were incubated for 600 sec before adding FCCP. Readings continued for 1000 sec. All conditions **(A, C, E)** were run in the same plate each of the three repeats and shared the same 0 µM control. Error bars are S.E.M. (n=3). ** *p*<0.01, *** *p*<0.001, \*\*\*\**p*<0.0001 (One way ANOVA for multiple comparisons).

The increase in AO fluorescence due to La^3+^ and Gd^3+^ was unexpected. A block in acidification typically manifests in a lack in AO fluorescence change. What would cause AO to increase in fluorescence? We hypothesize that this is linked to the effects of lanthanides on Ca^2+^-ATPases and Ca^2+^ channels. The yeast vacuole harbors the Ca^2+^-ATPase pump Pmc1 and the TRP ortholog Yvc1 [64,65]. Importantly, PMC family members as well as some TRP families can be inhibited by lanthanides [59,60,66]. In yeast Yvc1 releases Ca^2+^ from the vacuole lumen during osmotic shock, however Yvc1 is unlikely involved here because it is not active under the conditions of the assay [15,67–69]. If La^3+^ and Gd^3+^ inhibited Pmc1 (the primary Ca^2+^ uptake transporter on the yeast vacuole), the extraluminal Ca^2+^ would enter through the Ca^2+^/H^+^ antiporter Vcx1 [70]. Uptake through Vcx1 would result in the release of H^+^ from the vacuole lumen. Alternatively, the lack of Ca^2+^ gradient could inhibit V-ATPase function. In either case, H^+^ would not accumulate in the vacuole lumen. The ionic strength of La^3+^ and Gd^3+^ could keep AO from entering the vacuole that could lead to AO protonation (AOH). Protonation of AO would prevent its entry into the vacuole lumen where it would normally dimerize ([AOH]_2_) and fluoresce at 680 nm. Instead, the equilibrium of extraluminal AOH would shift to the monomeric form leading to the observed increased fluorescence at 530 nm. Clearly, these results are only the beginning a new set of experiments that are beyond the scope of a methods paper. Thus, the hypothesis is only a discussion point at this time.

## 4. Conclusion

Here we show the adaptation of a large volume single sample assay of vacuole acidification to a high-throughput microtiter plate format. Classically, acidification was show by shifts in absorbance of AO or fluorescence quenching of ACMA (9-amino-6-chloro-2-methoxyacridine) in cuvettes holding as much as 3 ml per reaction [34,50,71–73]. Cuvette based assays limit the amounts of conditions that can be tested with a single organelle preparation. We have scaled this assay down to 30 or 60 µl to fit in 384-well or 96 half-volume microtiter plates, respectively. This format allows for the screening of numerous reagents and conditions in a single round, which is of particular importance if testing low abundance and expensive reagents such antibodies. This also allows for the side-by-side comparison of genetic mutations and their wild-type parent.

Like many assays, there are limitations to what can be tested with this format. For instance, there is a maximum amount of DMSO and other organic solvents that can be used without damaging the vacuoles. As such, we have been unable to test specific small molecule inhibitors due to solvent effects. Other limitations can come from interfering with the reporter itself. Some small molecule inhibitors quench AO fluorescence in the absence of vacuoles. Thus, it is important to make sure that the reagents being tested do not indirectly interfere with membrane integrity or the reporter system.

## Abbreviations

AO: acridine orange
FCCP: Carbonyl cyanide-4-(trifluoromethoxy) phenylhydrazone
TRP: Transient Receptor Potential

## Author Contribution

C.Z., A.B., R.A.F. conceptualization, C.Z., A.B. Y.F. J.D.C. data curation; C.Z., A.B., R.A.F. formal analysis; C.Z., R.A.F. writing original draft; C.Z., R.A.F. writing rereview and editing; R.A.F. funding acquisition; R.A.F. supervision; R.A.F. project administration; R.A.F. resources.

## Acknowledgments

This research was supported by grants from the National Science Foundation (MCB 1818310, MCB 2216742) to RAF. J.D.C. was partially supported by an NIGMS-NIH Chemistry-Biology Interface Training Grant (5T32-GM070421).

